# Proteomic analysis of the Aggregation Factor from the sponge *Clathria (Microciona) prolifera* suggests an ancient protein domain toolkit for allorecognition in animals

**DOI:** 10.1101/2024.04.19.590289

**Authors:** Fabian Ruperti, Monika Dzieciatkowska, M. Sabrina Pankey, Cedric S. Asensio, Dario Anselmetti, Xavier Fernàndez-Busquets, Scott A. Nichols

## Abstract

The discovery that sponges (Porifera) can fully regenerate from aggregates of dissociated cells launched them as one of the earliest experimental models for cell adhesion and allorecognition studies in animals. This process depends on an extracellular glycoprotein complex called the Aggregation Factor (AF). However, our understanding of how animal adhesion and allorecognition mechanisms first evolved is complicated by the fact that the known components of the AF are thought to be unique to sponges. We used label-free quantitative proteomics to identify additional AF components and interacting proteins in the classical model *Clathria prolifera* and compare them to proteins involved in cell interactions in Bilateria. Our results confirm MAFp3/p4 as the primary components of the AF, but implicate related proteins with calx-beta and wreath domains as additional components. Using AlphaFold, we unveiled close structural similarities of AF components to distant homologs in other animals, previously masked by the stark decay of sequence similarity. The wreath domain, believed to be unique to the AF, was predicted to contain a central beta-sandwich of the same organization as the vWFD domain in extracellular, gel-forming glycoproteins in other animals. Additionally, we co-purified candidate AF-interacting proteins that share a conserved C-terminus, containing divergent Ig-like and Fn3 domains, a combination also known from IgCAMs. One of these, MAFAP1, may function to link the AF to the surface of cells. Our results highlight the existence of an ancient toolkit of conserved protein domains regulating cell-cell and cell-ECM interactions in all animals, and likely reflect a common origin of cell-adhesion and allorecognition.

## Introduction

The theoretical requirements for the evolution of obligate multicellularity include adhesion mechanisms for cell-cell attachment (1), signaling mechanisms to guide development and enable coordinated interactions between cells (2), and allorecognition mechanisms to distinguish self from nonself (3). In traditional bilaterian research models, much is known about how these theoretical requirements are met. But comparative studies of non-bilaterian animals such as sponges, ctenophores, and cnidarians are needed to reconstruct the earliest events in the evolution of animal multicellularity.

In 1907, Henry Wilson’s seminal experiments with *Clathria (Microciona) prolifera* showed that sponges can be dissociated into a heterogeneous suspension of cells which - under the right conditions - reassemble into a functional, intact organism (4). This process is highly specific and depends on the selective attachment between cells of the same genotype (5). For over a century since, researchers have used ‘aggregation assays’ to study the mechanisms of cell adhesion and allorecognition in sponges. The prevailing model proposes that species-specific aggregation depends on a sulfated proteoglycan termed the ‘Aggregation Factor’ (hereafter, AF) found in the extracellular matrix and bound to the surface of cells (6– 12). In aggregation assays performed with synthetic beads coupled to AF molecules from different species (13, 14), the beads sorted out and associated only with those carrying AFs from the same species, showing that AFs are sufficient for species-specific aggregation *in vitro*.

The composition of the AF varies between species, consisting of 50-70% protein and 30-50% carbohydrate (15–17). In some species, individual AF molecules can be linear, similar to proteoglycans in other animals, but others have a unique ‘sunburst’ architecture with a central ring and radiating arms (15, 17) (**Figure 1A**).The well-studied sunburst-like AF of *C. prolifera* has a molecular weight of *∼* 2 x 10^7^ Da and is composed of two main proteins that derive from the precursor protein MAFp3/p4 (**Figure 1B, 3A**) (15, 18–20). Twenty copies of the *∼* 50 kDa MAFp3 subunit comprise the inner ‘ring’ of the sunburst, and each is decorated with a 200 kDa glycan (g-200) (21, 22). Twenty subunits of the *∼* 400 kDa MAFp4 subunit comprise the radiating ‘arms’ of the AF, each of which is attached to the MAFp3-ring and decorated with about 50 copies of a 6 kDa glycan (g-6) (15, 18–20, 23). The differences of the glycoconjugates to known glycosaminoglycans prompted the name ‘glyconectins’ (24, 25).

**Figure 1.**
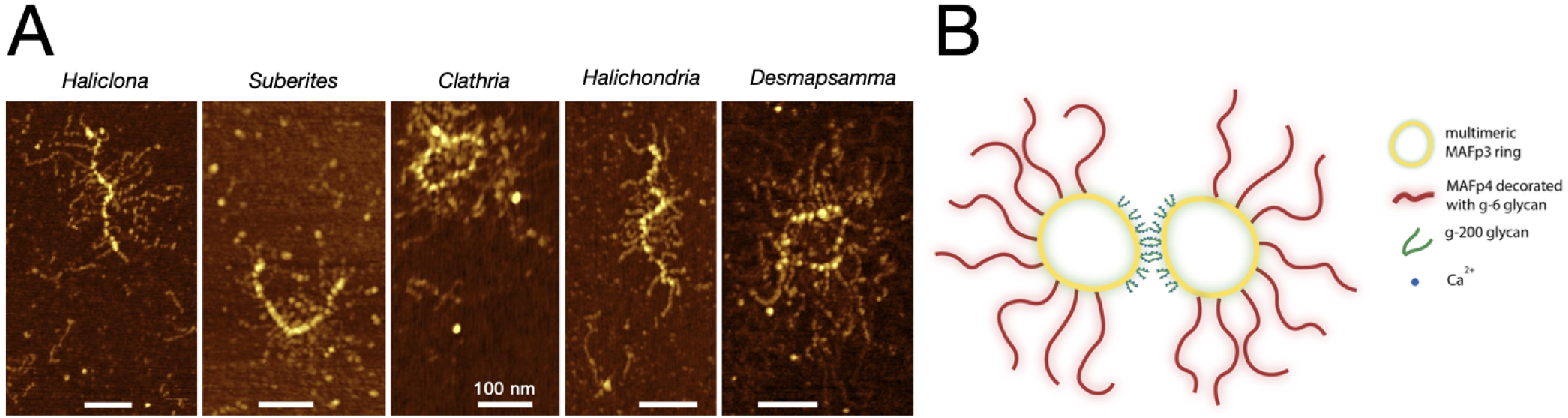
The aggregation factor is composed of glycoproteins that form linear or circular structures with radiating arms. **(A)** Atomic force microscopy (AFM) images of linear versus ring-like AF purified from different sponge species. AF purification and AFM imaging were performed as previously described (29–31) **(B)** Cartoon depiction of the known components of *C prolifera* AF. Wreath-domain containing MAFp3 makes up the central ring and is decorated with g-200 glycans responsible for Ca^2+^-dependent AF-AF interactions. Calx-beta domain-containing MAFp4 arms are decorated with g-6 glycan responsible for Ca^2+^-independent AF-cell interactions.

Ca^2+^-dependent carbohydrate-carbohydrate interactions between g-200 glycans determine the specificity needed for AF-AF binding (21) (**Figure 1B**). In contrast, AF-cell binding is mediated by Ca^2+^-independent interactions between the g-6 glycan with an unidentified 68 kDa lectin-like protein that is hypothesized to then bridge to an integral membrane receptor (20, 26). In support of this, AF–AF and AF-cell binding affinities were recovered upon chemical crosslinking of protein-free glycans isolated from the AF complex into large multivalent structures (22, 23, 27). Also, glycan-coated beads aggregate according to their species of origin, and live cells selectively bind to species-specific, glycan-coated surfaces (28).

The *C. prolifera* AF has been described to contain at least 11 different protein subunits that could be dissociated upon Ca^2+^ removal (32). However, MAFp3 and MAFp4 are the only unequivocally identified components and contain at least 15 calx-beta domains and a novel C-terminal structure termed the “wreath” domain (33). Calx-beta domains typically bind Ca^2+^ and in the extracellular matrix are known only from the human Fras1-related extracellular matrix protein (FREM-1), adopting an Ig-like beta-sandwich fold, similar to Ig-like and Fn3 domains (SCOP family 48725).

The predicted wreath domain in the MAFp3 subunit appears to be restricted to demosponges – even to the exclusion of other sponge lineages such as calcareans, homoscleromorphs, and hexactinellids (33). This limited phylogenetic distribution has been interpreted as evidence that the demosponge AF is unrelated to adhesion and allorecognition mechanisms in other organisms, or that rapid evolution has erased the traces of homology between allorecognition systems in divergent animal lineages.

Comparative genomics data support the possible existence of additional AF protein components. Specifically, demosponge genomes encode many additional genes with homology to MAFp3/p4. Some have a conserved wreath domain in combination with various other structural elements including calx-beta, von Willebrand Factor A/D (vWFA/D), and Plexin-Semaphorin-Integrin (PSI) domains. Lower confidence candidate AF components lack a wreath domain but have conserved calx-beta domains and BLAST to MAFp3/p4 (33). These have not been experimentally validated to interact with the AF, but in the genome of *Amphimedon queenslandica* they are found in syntenic clusters and exhibit high transcript polymorphism – common characteristics of allorecognition genes.

Other AF or AF-interacting proteins have also been detected experimentally but remain incompletely identified. Specifically, Varner (12, 34) used the AF as a probe to purify candidate AF-binding proteins in *C. prolifera*. Two proteins of *∼* 68 kDa and *∼* 210 kDa were found to bind with high affinity to the AF in the extracellular matrix, and to bind with high affinity to the cell surface (12, 26, 34). Also, using dissociative gel fractionation of the *C. prolifera* AF, Fernandez-Busquets and colleagues (19) identified two discrete bands at 210 kDa (p210) and 2,000 kDa (S2). They determined the N-terminal peptide sequence from these bands, but protein database searches at the time did not yield significant similarities to known proteins. They speculated that the 210 kDa band was likely the same as that identified by Varner (12, 34), and demonstrated that it was a glycosylated protein exhibiting interindividual polymorphism in the glycan moiety (18). This observation was consistent with the role of AFs as determinants of individuality within a species and suggests that differential glycosylation could be a way to distinguish between self and nonself (30).

To comprehend how the AF-model of adhesion/allorecognition fits into the broader context of multicellular evolution, a deeper understanding of its protein backbone is required. Here, we used a proteomics approach to identify proteins associated with the AF in *C. prolifera*. Consistent with prior studies, we found that the most abundant protein components of the AF were MAFp3/p4. Other abundant proteins in purified AF samples fell into one of two major categories: 1) like MAFp3/p4 they contain a predicted wreath domain and/or calx-beta domains, or 2) they have a C-terminus composed of Ig- and Fn3-like domains that potentially serves as an interaction interface with the AF. All have low predicted isoelectric points, consistent with having a negative charge at neutral pH – a common characteristic of secreted Ca^2+^-binding proteins in Bilateria. In contrast to sequence-based comparisons, structural analyses indicate that the core beta-sandwich of the AF wreath domain is nearly identical to vWFD domains present in gel-forming extracellular glycoproteins of bilaterians. Likewise, the C-terminus of candidate AF-interacting proteins is found to have extensive structural similarity to IgCAMs such as neural cell adhesion molecules (NCAMs). These results expand our understanding of the structure, composition and endogenous interactions of the AF, and provide clues to the existence of an ancient protein domain toolkit for cell-ECM interactions that is shared between sponges and bilaterians.

## Results

### *De novo* assembly of a *Clathria prolifera* proteome database

To create a reference database for proteomic analyses, we first sequenced and assembled the transcriptome of *Clathria prolifera* using RNA isolated from whole, adult tissues. A Novaseq pe150 run (Novagene) produced 42,531,984 x 150 bp paired-end reads, which were assembled to yield 22,794 predicted transcripts. The final assembly had a BUSCO v3 completeness score of 89.6% and a BUSCO v4 completeness score of 85.1%. We deposited raw reads in the NCBI Sequence Read Archive (SRX18275041) and archived the TransPI assembly and corresponding peptide predictions on Figshare (35).

We prepared Aggregation Factor (AF) samples for proteomic analyses in three ways (**Supplemental Figure 1**). First, we isolated a ‘Crude’ AF extract based on the method of Humphreys (36). Briefly, we dissociated live tissue in Ca^2+^/Mg^2+^-free seawater (CMFSW), removed cells and spicules by centrifugation, and then precipitated the AF from the supernatant by the addition of CaCl_2_ to form a red, gel-like pellet. To remove pigments, membranes, and possible other contaminants, we re-dissolved Crude AF in CMFSW, centrifuged it, and passed it through a 0.22 µm filter (see Varner (34)). When we again added CaCl_2_ to precipitate the AF it formed a fluffy, white pellet (‘Filtered’ sample). As a final purification step we re-solubilized the precipitate from the filtered sample for fractionation by size exclusion chromatography on an s500 column (Cytiva). The upper limit of the fractionation range of this column is 2 x 10^7^ Da, which is the reported molecular weight of the *C. prolifera* AF (15, 18, 19). We then combined and concentrated the fractions expected to contain the AF for analysis as the ‘SEC’ sample.

Proteomic analyses of the Crude AF extracts detected a total of 690 predicted proteins. Analysis of the ‘Filtered’ and SEC samples resulted in the detection of 437 and 696 proteins, respectively. A total of 319 unique proteins were common to all samples (**Figure 2A**). The relative proportion of the proteins within each sample was calculated by dividing the protein intensity (based on MS1 label-free quantification of unique+razor peptides) by the sum of all peptide intensities from each sample. Most detected proteins had very low relative abundance (**Figure 2B**). Proteins detected exclusively in any one of the three samples showed only a proportion < 5% in their respective samples.

**Figure 2.**
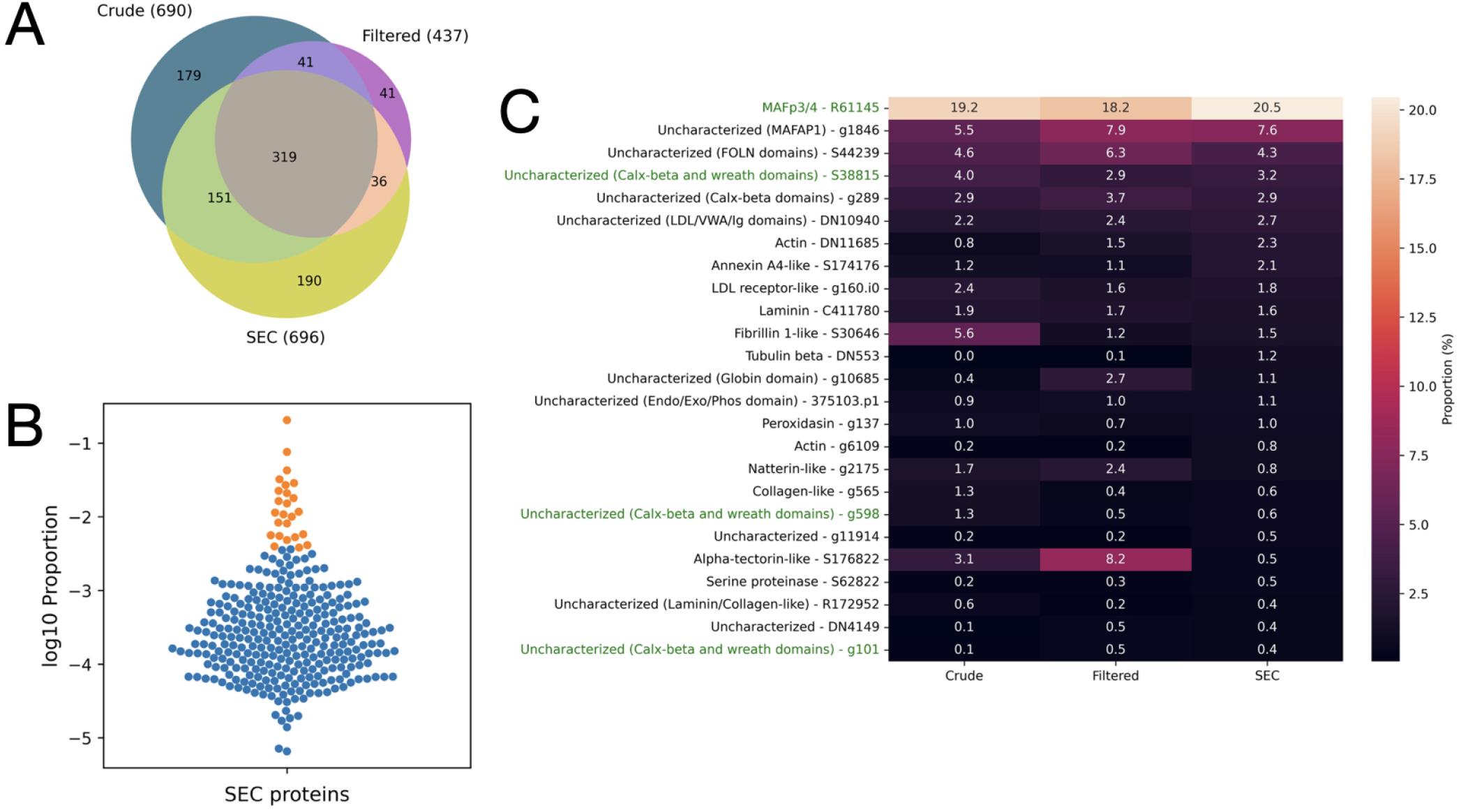
Protein abundance in each LC-MS/MS sample. **(A)** Venn diagram of proteins detected in each of the Crude, Filtered, and SEC samples. **(B)** Violin plot showing protein proportion in the SEC sample of the 319 proteins found in all samples plotted on a log10 scale. The top 25 most abundant proteins are highlighted in orange. **(C)** The 25 proteins with highest relative proportion in each of the Crude, Filtered, and SEC samples. There was high concordance between samples, with the exception of the S176822, which had low abundance after size exclusion chromatography in the presence of EDTA. Proteins containing a wreath domain are highlighted in green.

### AF samples contain MAFp3/p4 and related wreath-domain proteins

The high diversity of proteins detected in all samples indicate that our methods did not produce a highly purified preparation of the AF, or that high levels of glycosylation of the AF interfered with analysis by LC-MS/MS leading to an apparent overrepresentation of low-level contaminants. Still, we reasoned that the best candidate AF components and/or binding partners were those that were highly represented in all samples (**Figure 2B**). Indeed, the most abundant proteins detected in all samples were the known AF components MAFp3 and MAFp4, which together comprised 19.2% of the Crude sample on average, 18.2% of the Filtered sample, and 20.5% of the SEC sample (**Figure 2C**)

The transcriptome assembly was found to encode a *∼* 430 kDa MAFp3/p4 precursor protein (a prediction that does not take into account glycosylation). The MAFp4 region has 29 calx-beta domains (**Figure 3A**), and as previously described, the MAFp3 region contains the wreath domain. In addition to MAFp3/p4 we also detected an additional eight wreath domain-containing proteins in the *C. prolifera* transcriptome assembly (**Supplemental File 1**). Of these, three (**Figure 3A**) were also found within the top 25 most abundant proteins in our proteomics datasets (with S38815 being the fourth most abundant protein in the SEC sample) (**Figure 2C**) and each was predicted by Interpro (37) to have between 8 and 29 N-terminal calx-beta domains.

**Figure 3.**
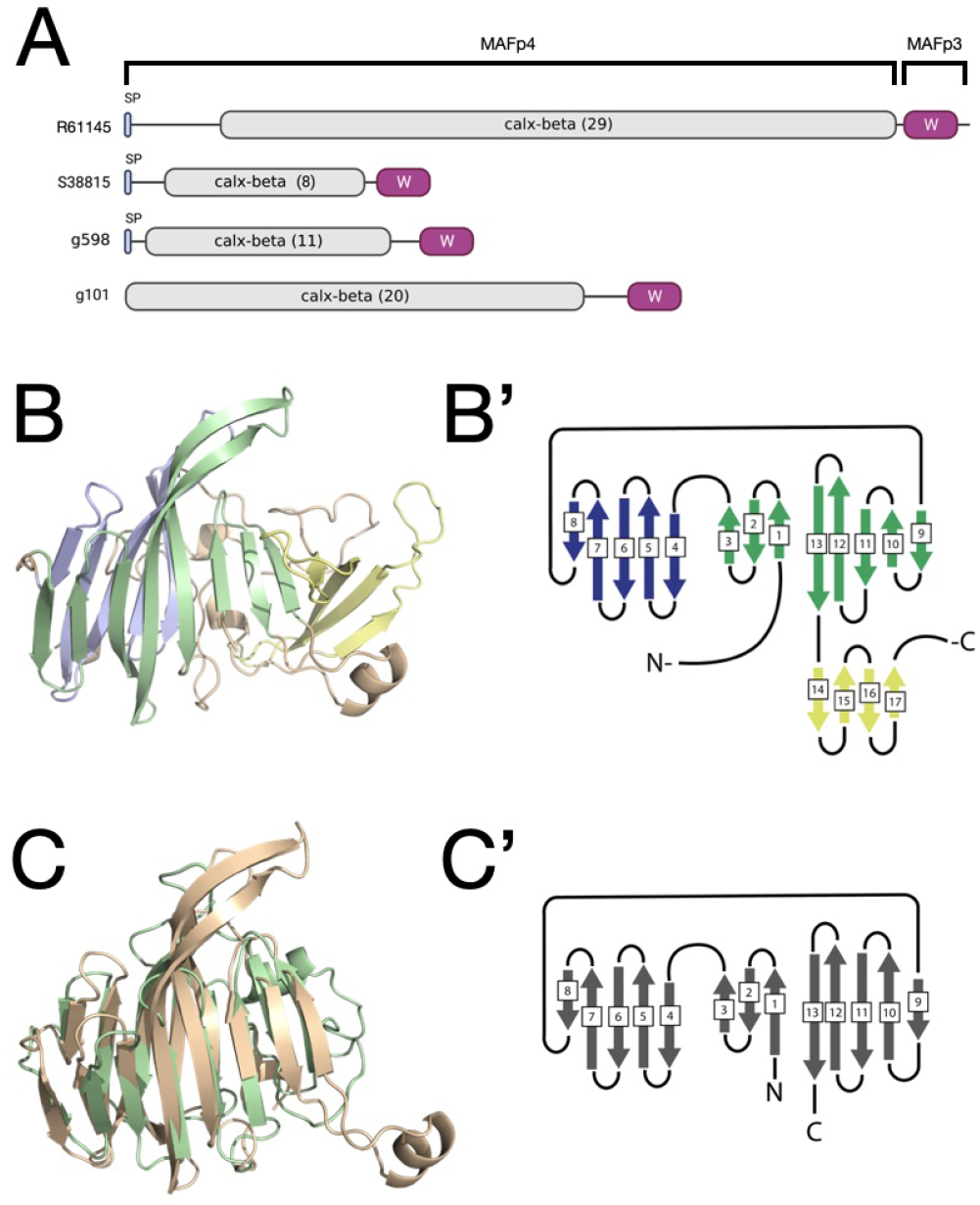
Wreath domain-containing proteins detected in AF proteomic samples have structural similarity to the vWFD domain. **(A)** Wreath domain-containing proteins in order of their relative proportion in AF proteomic datasets. Proteins are highlighted in green in Figure 3C. **(B)** Colabfold prediction of MAFp3/p4 wreath domain, with beta-strands highlighted to reflect their position in the tertiary structure. Disordered N- and C-termini were manually trimmed for visibility. **(C)** Superposition of MAFp3/4 wreath domain beta-sandwich (beige) with VWFD1 domain of *H. sapiens* Mucin-5AC (UniprotID: P98088; green). RMSD = 2.22 over 481 atoms. Beta strand switch 1-3/4-8/9-13 between sandwiches is conserved in VWFD domains (40).

### The wreath-domain is structurally similar to vWFD domain

The wreath domain has previously only been detected based upon multiple sequence alignment of MAFp3 homologs (33). We used Colabfold (38) to predict the wreath domain structure (average pLDDT = 72.2) (**Supplemental File 2**). The model indicates an essentially all-beta structure with isolated short alpha-helices in the periphery. A long, unfolded C-terminal domain is predicted to be disordered (prediction by IUPred3) (39). A central beta-sandwich is formed by two twisted anti-parallel beta sheets, containing 8 (green, strand 1-3, 9-13) and 5 (blue, strand 4-8) beta strands of varying lengths (**Figure 3B**). The beta-sheets are connected by strands 3 and 4 as well as 8 and 9. Additionally, the structure has a curved, anti-parallel beta sheet (4 beta strands, 14-17) adjacent to the beta-sandwich (yellow).

Since hidden-markov-model (HMM)-based searches for wreath domain containing proteins in other species indicate that this domain is demosponge-specific (33), we instead searched for structurally similar proteins using the Colabfold prediction for the MAFp3 wreath domain in Foldseek, as this method should find structurally similar proteins irrespective of their sequence conservation (38, 41, 42). Top hits were to wreath domain-containing proteins in *Clathria prolifera* and other demosponges included in the AlphaFold DB, such as *Suberites domuncula* and *Amphimedon queens-landica*. However, a structurally similar protein (UniProt A0A1X1QQN9) was also detected from the bacterial species *Cycloclasticus sp. M*. (43). Unexpectedly, this sequence had 82% sequence identity to the *C. prolifera* MAFp3/4 wreath domain (**Supplemental Figure 2**), albeit only part of the beta-sandwich region is conserved. This bacterium was isolated as a symbiont of a poecilosclerid sponge, suggesting horizontal gene transfer from the sponge to the bacterium.

Notably, using the central beta-sandwich motif of the wreath domain as a query for Foldseek, we also detected hits in bilaterians, including proteins such as alpha-tectorin, mucin, IgGFc-binding protein or otogelin. A common feature of these proteins is the presence of a von Willebrand Factor D (vWFD) domain. Exemplary structural superposition of the wreath domain with the first vWFD domain of human mucin 5AC highlighted a highly conserved beta-sandwich, showing identical position switch of beta strands between the two sheets of the sandwich (40) (**Figure 3C**), despite an overall low sequence identity of 12.4%, which usually precludes detection by sequence-based similarity searches. Interestingly, the vWFD domain is found in secreted, gel-forming glycoproteins (44) and is pivotal for the platelet aggregation function of the name-giving von Willebrand Factor (45), in line with the function of wreath domain-containing AF proteins.

### Aggregation factor-associated proteins share a conserved C-terminus composed of divergent Immunoglobulin and Fibronectin 3-related domains

The second most abundant protein in the SEC dataset (and in all datasets when combined) corresponded to transcript g1846, and is hereafter referred to as MAF-associated protein 1 (MAFAP1). In the Filtered sample, MAFAP1 was slightly less abundant (7.9%) than an uncharacterized NIDO and calx-beta domain-containing protein (8.2%), but this protein only showed a proportion of 0.5% in the SEC sample, indicating its effective separation from the AF by column chromatography. This may indicate that it is either not stably associated with the AF, or that this association was disrupted by short-term exposure to EDTA (which was used to solubilize the Filtered sample for column chromatography).

MAFAP1 is predicted as an *∼* 100 kDa protein with a signal peptide, two N-terminal immunoglobulin-related (Ig) domains, and an intrinsically disordered region composed of 69 pentapeptide repeats with the consensus amino acid sequence PETDA (**Figure 4A**). Notably, it contains the peptide sequence **ELIDYETFSDGRVL** identified from the 210 kDa AF component purified by Fernàndez-Busquets and collegues (19), but which could not be further resolved at the amino acid level at the time. BLAST searches using MAFAP1 as a query against the NCBI non-redundant protein database and against the more inclusive Eukprot3 database (46) resulted in low-confidence hits to proteins with repeat regions that are only superficially similar to the disordered PETDA repeat region. No candidate MAFAP1 homologs were detected by BLAST search in any other species, including in other sponges.

**Figure 4.**
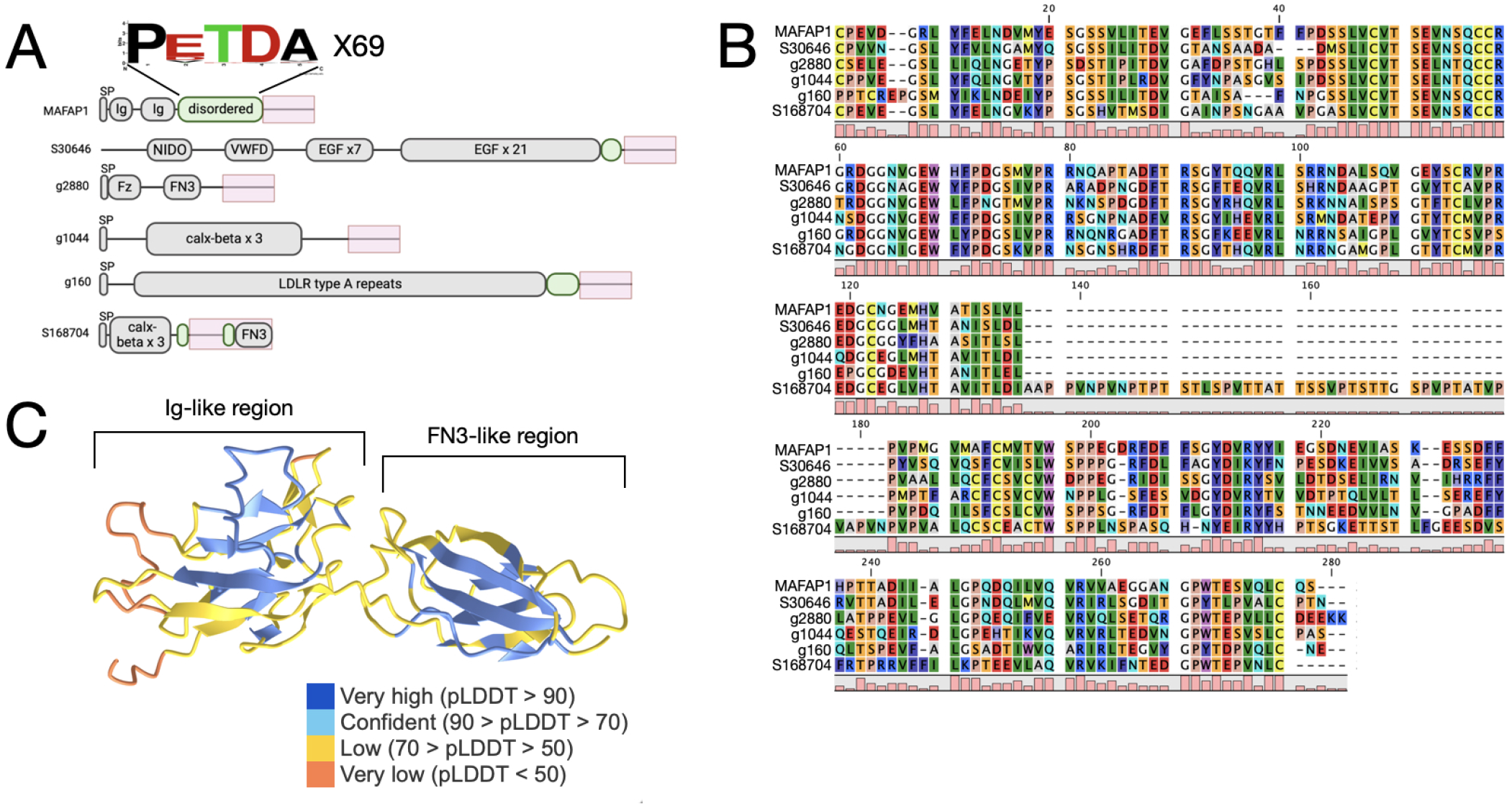
Top MAFAP1 hits in the predicted proteome are unrelated but share a highly conserved, candidate AF-interaction domain. **(A)** Domain architecture of *C. prolifera* proteins that contain a conserved AF-interacting region (pink box) in common with MAFAP1. Like MAFAP1, three of these also contain a disordered repeat region (green oval) adjacent to the AF-interacting region. All but S168704 were detected in proteomics results for the AF. **(B)** Multiple sequence alignment of the AF-interacting regions from proteins depicted in panel A. **(C)** Colabfold prediction for the structure of the AF-interacting region, highlighting a divergent Ig-like domain (region 1) and a divergent Fn3-like domain (region 2).

We next searched by BLAST for additional MAFAP1-related proteins in the *C. prolifera* transcriptome. Again, there were no candidate homologs, but 5 proteins had high identity-score matches to a 226 aa region of the C-terminus (**Figure 4A,B**). These hits were otherwise distinct from MAFAP1 and from each other, and included a fibrillin-like protein (S30646, containing NIDO, vWFD and EGF domains), a secreted frizzled homolog (g2880, containing Fz and FN3 domains), a MAFp3/p4-related protein with calx-beta domains but lacking a wreath-domain (g1044), a low-density lipoprotein receptor-related protein (g160), and an uncharacterized protein with calx-beta and Fn3 domains (S168704).

All but S168704 (which has the least-conserved C-terminal sequence, including an unstructured, repetitive insert sequence) were present in the proteomics datasets, and of these the fibrillin-like protein (S30646) was among the top 25 most abundant proteins detected overall (**Figure 2C**). This suggested that the shared C-terminal region may represent an important binding interface for interactions with the AF. Hereafter, we refer to this conserved region as a candidate ‘AF-interacting region’.

Structural analysis of the candidate AF-interacting region by Colabfold indicated the presence of two discrete elements composed of beta-sheets and similar to immunoglobulin (Ig-like) (region 1) and fibronectin 3 domains (Fn3) (region 2) (**Figure 4C**) (**Supplemental File 3**). Indeed, the top 5 Fold-seek search results against AFDB-Swissprot using the predicted structure of MAFAP1 C-terminal domain as a query, were neural cell adhesion molecules (NCAM) from various bilaterians, mapping to Ig-like and Fn3 domains, despite a sequence identity < 15%. Structural superposition and sequence alignment of region 1 to NCAM Ig-like-domain supports the close structural similarity and highlights the conservation of two critical cysteine residues that form a stabilizing disulfide bridge in Ig-domains (47) (**Figure 5A, A’**). Likewise, region 2 of the MAFAP1 C-terminal domain is structurally very similar to the adjacent Fn3 domain of the respective NCAM proteins (**Figure 5B, B’**).

**Figure 5.**
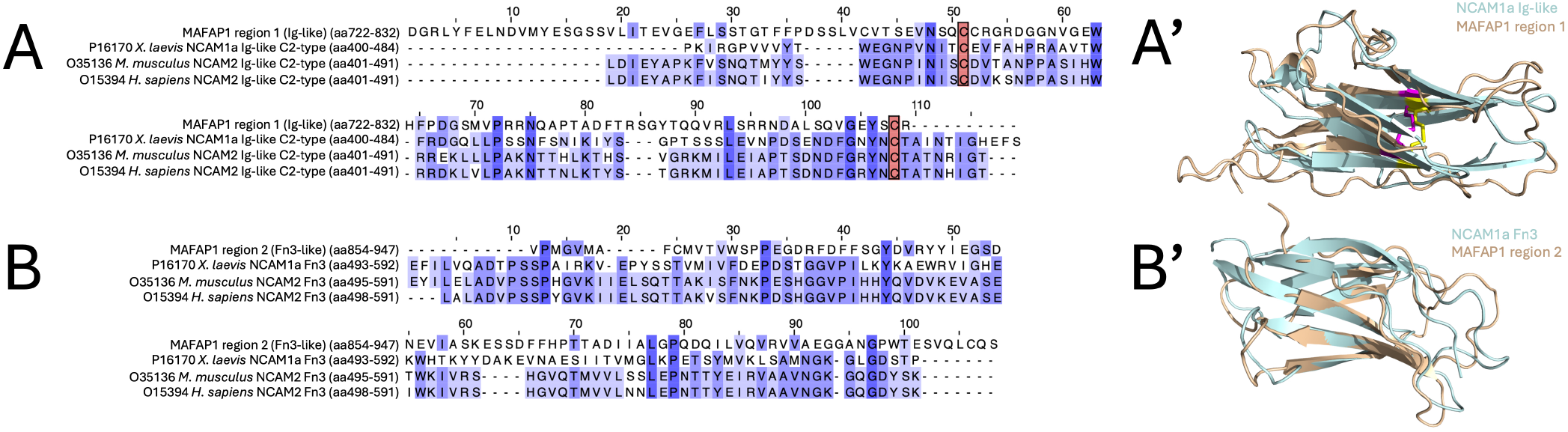
Sequence and structural alignment of MAFAP1 C-terminal domain. **(A, A’)** Sequence alignment of region 1 of MAFAP1 C-terminal domain to NCAM Ig-like-domains (C2-type) from top Foldseek hits. Conserved cysteine residues, typical for Ig-like domains, are highlighted in red. Structural superposition of Colabfold prediction of region 1 of MAFAP1 C-terminal domain (beige, aa722-853) to NCAM1a Ig-like-domain (UniprotID: P16170, aa400-484, blue) (RMSD: 4.3 over 80 residues). Conserved cysteine residues that form a stabilizing, intramolecular disulfide bridge are highlighted in magenta (for MAFAP1) and yellow (for NCAM1a). **(B, B’)** Sequence alignment of region 2 of MAFAP1 C-terminal domain to NCAM Fn3-domains from top Foldseek hits. Structural superposition of Colabfold prediction of region 2 of MAFAP1 C-terminal domain (beige, aa854-947) to NCAM1a Fn3-domain (UniprotID: P16170, aa493-592, blue) (RMSD: 2.9 over 88 residues).

NCAM belongs to the Immunoglobulin Superfamily Cell Adhesion Molecules (IgCAMs or IgSF CAMs) (48) which regulate cell-cell/ECM adhesion via extracellular Ig-like domains and are anchored via transmembrane (TM) regions or GPI. Although none of the proteins in *C. prolifera* with AF-interacting region are predicted to contain a TM region or GPI-anchor (prediction via TMHMM and NetGPI; (49, 50)), all are secreted extracellular proteins that presumably are able to interact with either the glycan or protein component of the AF.

### Proteins with conserved AF-interacting regions in other sponge species

MAFp3/p4 and other wreath domain-containing proteins are widely conserved in demosponges. To search for proteins that contain the putative AF-interacting region in other sponge species we developed a HMM from the alignment of the AF-interacting region of *C. prolifera* proteins (**Supplemental File 4**). We then searched the predicted proteomes of *Ephydatia muelleri* (51), *Tethya wilhelma* (52), *Amphimedon queenslandica* (53) (all heteroscleromorph demosponges), and *Cladhorizida sp*. (54); a poecilosclerid demosponge, like *C. prolifera*). None were found to have proteins with a conserved AF-interacting region.

When we expanded this search to include partial transcriptome data available from additional poecilosclerid sponges, we found evidence for highly conserved AF-interacting regions in proteins from *Tedania anhelans* (55), *Crella elegans* (56), and *Phorbas areolata* (57). In all but two sequences from *Tedania anhelans*, the transcriptome assemblies were too fragmentary to characterize features of these proteins beyond the presence of the conserved AF-interacting region. But, like *C. prolifera* proteins that contain the AF-interacting region, two *T. anhelans* sequences were found to encode a CRDFz domain, and one was found to have calx-beta domains (**Supplemental File 5**). Using BLAST, the top hits of these proteins were secreted frizzled-related proteins and frizzled receptors in other sponges, and to MAFp3/p4, respectively.

### Other abundant AF-associated proteins contain known domains of extracellular proteins

Four additional proteins were relatively abundant in all proteomic datasets (**Figure 2C**). These included a highly conserved annexin A4-like protein (found in collagen-containing extracellular matrix of vertebrates) and three uncharacterized proteins. One of those (S44239) contains four follistatin-N-terminal domain-like domains which are also found in the extracellular matrix proteins Agrin and SPARC. The second uncharacterized protein (g289) again contains calx-beta domains, as detected in MAFp4 and other AF-associated proteins. The third uncharacterized protein (DN10940) has a signal peptide, LDL, Ig, and von Willebrand Factor A domains (vWFA), and similarity by BLAST search to contactin, neurofascin, and NCAM – presumably due to the presence of Ig-like domains. The Ig/vWFA region is predicted by Interpro to relate to the Basigin family, which includes two protein subfamilies: neuroplastin and basigin. Both have extracellular Ig-like domains and are glycosylated. Neuroplastin functions in neuronal cell adhesion, whereas basigin has more diverse functions including stimulating the production of matrix metalloproteinases in fibroblasts (58). Among the top 25 proteins we furthermore identified proteins such as laminin, which add to the pool of secreted glycoproteins that regulate cell-cell and cell-ECM interactions.

### AF proteins share a low predicted isoelectric point (pI)

The Glu and Asp residues of the pentapeptide repeats of MAFAP1 lead to an exceptionally low predicted isoelectric point (pI) of 3.56 (prediction with the python implementation of ‘Peptides’ (https://pypi.org/project/peptides/)) 59). Proteome-wide comparison of predicted pIs, Glu and Asp proportions reveal that putative AF components share a significantly lower pI (4.4 *±* 1.4 for top 25 AF proteins vs 7.1 *±* 2.1 for whole proteome) (**Supplemental Figure 3A**) due to high proportions of Glu (7.5 *±* 1.8%) and Asp (7.7 *±* 2.1%) residues. Comparison to amino acid compositions from AF preparation of the related sponge *Clathria (Microciona) parthena* (15) supported our results and suggested that Henkart and colleagues likely worked with a pure MAFp3/p4 and MAFAP1 sample due to the high proportion of Glu and Asp residues (**Supplemental Figure 3B**). In neutral extra-cellular environments, AF protein components are therefore predicted to be negatively charged, favoring Ca^2+^-binding.

## Discussion

Studies of the AF in sponges have long emphasized its role in providing the adhesive force and specificity needed for cell aggregation (29). However, as the details of its molecular composition unfolded, studies in other animal models led to the discovery of integral membrane receptors such as cadherins (60, 61) and integrins (62, 63) at specialized cell junctions. These molecular differences, together with the apparent lack of cell junctions detectable in sponges by transmission electron microscopy, contributed to a narrative that sponges lack ‘true tissues’ and are instead organized as more of a transient collection of pluripotent cells embedded in a common AF-rich extracellular matrix (64, 65). However, from comparative genomics we have since learned that, in addition to the AF, sponges have homologs of cell junction adhesion receptors and their associated effectors such as catenins, talin, and vinculin (53, 66–68). Moreover, some of these proteins have been shown to interact and localize to specialized junctions in sponges, similar to epithelial tissues in other animals (69–71). These discoveries raise new questions about whether (and how) the AF may interact with cell junction proteins in the context of adhesion and allorecognition.

A challenge for integrating our new understanding of sponge cell junctions with the AF model of adhesion and allorecognition is that the former have been exclusively studied in the context of intact tissues, whereas the latter has largely been studied in the context of aggregation assays. However, it has been hypothesized that AF receptors may interact with integrins (64), which is interesting in light of the fact that some integrins have calx-beta domains (72) like many candidate AF components and AF-interacting proteins. Also, sponge integrins have unexpectedly been shown to localize to cell-cell contacts in addition to cell-ECM contacts (71), which is consistent with an adhesion model where cells are linked together through the AF as an extracellular intermediary.

The goal of the current study was to apply proteomic methods to identify protein components of the AF together with interacting proteins that may give new clues to its evolution and relation to sponge cell junctions. The detection of MAFp3/p4 as the most abundant proteins in our purified AF samples validates the efficacy of this approach. Also, our results support the prediction from comparative genomic analyses that additional calx-beta and wreath-domain containing proteins may also be components of the AF (33).

The strongest clue linking the AF to extracellular matrix components in other organisms comes from the predicted structure of the wreath domain itself. The central beta-sandwich of the wreath domain is highly similar to vWFD domain and to the C-terminal domain of Repulsive Guidance Molecules (40), despite that its sequence is divergent beyond recognition from both. Moreover, just as the wreath domain of MAFp3 forms the ring-like core of the *C. prolifera* AF, in vitro expressed vWFD domains from mucins multimerize into ring structures (73) of the same size. These similarities suggest homology between the AF wreath domain with the vWFD domain, or else remarkable convergent evolution of a beta-sandwich that appears in secreted, heavily glycosylated proteins that function in cell-cell and cell-ECM interactions.

Our results also corroborate the 210 kDa protein independently detected by Varner (12) and Fernàndez-Busquets (19) (which we named MAFAP1) as a candidate AF-interacting protein. In our datasets, this was the 2nd most abundant protein overall. The protein sequence of MAFAP1 has a predicted MW of only *∼* 100 kDa, but this difference from previous studies possibly reflects the glycosylation state of the endogenous protein, or perhaps that it forms homodimers. Although no clear MAFAP1 homologs were detected in other (even closely related) species, sequence analysis did reveal a candidate C-terminal AF-interacting region shared with 5 otherwise unrelated *C. prolifera* proteins. Structural prediction and search with this region showed close similarity to Ig and Fn3 domains of proteins in Bilateria that function in cell-cell/ECM adhesion, including NCAM, a member of the IgCAM superfamily. With more structural predictions available for sponge proteomes in the future, there may also be more undetected matches to the AF-interacting region within distantly related species. Although the exact interaction between AF and the putative AF-interacting region is still unresolved, we hypothesize that MAFAP1 in particular could link the AF to cell surfaces.

This study highlights the existence of a conserved toolkit of protein domains that are present in proteins regulating cell-cell and cell-ECM adhesion and recognition, both in Bilateria and in sponges. These include vWFD, Ig-like, Fn3 and calx-beta domains, with the latter three adopting similar Ig-like beta-sandwich folds. Although the decay of protein sequence similarity often prohibits the assumption of homology, the advance of tools such as AlphaFold allowed us to trace back the existence of those domains in the sponge AF and AF-associated proteins. On top of conserved protein domains, physico-chemical parameters such as low predicted pIs, which in turn lead to negative charges at neutral (extracellular) pH, appear to be conserved in secreted, Ca^2+^-binding proteins such as osteopontin (pI = 3.5) (74, 75) or the cell adhesion protein fibronectin (predicted pI = 5.5). In the future, it will be interesting to apply the same unbiased proteomics approach that we used in this study to examine the AF composition and interactions in progressively distantly related demosponge species, which may provide additional clues to how the AF evolved and relates to adhesion and allorecognition mechanisms in other sponges and in non-sponge animals.

## ACKNOWLEDGEMENTS

We thank Bernadette Doyle for help in assembling the *C. prolifera* transcriptome and in purification of the AF, the Savitski team at EMBL for their help with proteomic data analysis, Detlev Arendt for the support to FR, and Ana Riesgo for sharing transcriptome assemblies of poecilosclerid sponges. This work was supported by a National Science Foundation grant (IOS:2015608) to SA Nichols.

## AUTHOR CONTRIBUTIONS

F.R. data analysis and writing, M.D. proteomic data collection and writing, S.P. data analysis and writing, C.A. SEC data collection, D.A. AFM data collection, X.F.-B. AFM data collection and writing, S.N. experimental design, data collection, data analysis and writing.

## Methods

### Data availability

The mass spectrometry proteomics will be deposited to the ProteomeXchange Consortium (http://proteomecentral.proteomexchange.org) via the PRIDE (76) partner repository upon final publication. Any additional information required to reanalyze the data reported in this paper is available from the lead contact upon request.

Raw *C. prolifera* teanscriptomic reads are deposited in the NCBI Sequence Read Archive under the identifier SRX18275041. The TransPi assembly and corresponding peptide predictions are archived on Figshare (35).

## Computational resources table

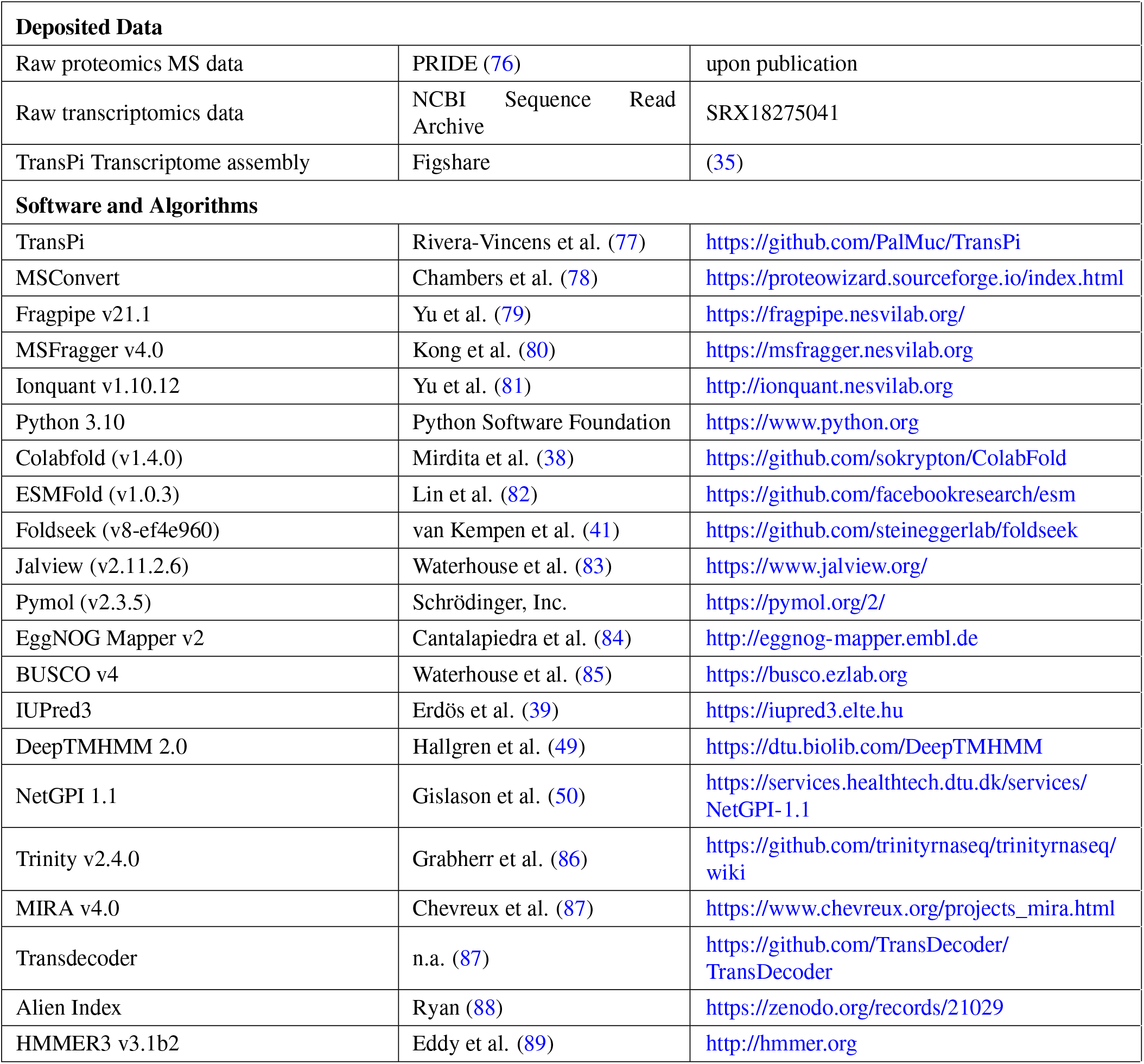

### Transcriptome sequencing and assembly

We acquired live samples of *C. prolifera* from the Marine Resources Center of the Marine Biological Laboratory at Woods Hole, cleaned tissues of debris and macroscopic contaminants, then ground them to a fine powder in liquid nitrogen. Using Trizol Reagent (Thermo Fisher Scientific), we isolated total RNA and shipped it to Novogene (Chula Vista, CA USA) for library preparation and Novaseq pe150 bp sequencing. We assembled and annotated the transcriptome using the TransPi pipeline (77) as implemented on the University of Denver High Performance Computing Cluster.

### Aggregation Factor Preparation

We purified crude AF using the (approximate) method of Humphrey (36). Briefly, we cut *∼* 100 g of tissue into 1 cm pieces, washed them in cold Calcium Magnesium Free Seawater (CMFSW; 2 mM NaHCO_3_, 462 mM NaCl, 7 mM Na_2_SO_4_, 10.7 mM KCl, pH 7.2) for 10 minutes, then rinsed them again briefly in CMFSW. We then squeezed tissue fragments through a fine synthetic mesh into a beaker containing 100 mL cold CMFSW, aliquoted dissociated cells into 50 mL conical tubes, and placed them on a rotator for 2 hrs at 4 ºC. We separated cells from the AF-containing supernatant by centrifugation in a swinging bucket rotor at 1,500 x g for 5 min at 4 ºC. To remove any remaining insoluble debris, we then centrifuged the supernatant at 10,000 x g for 15 min at 4 ºC. To precipitate the AF from the supernatant, we added CaCl_2_ to a final 20 mM concentration and placed the beaker on a stir plate at 4 ºC for *∼* 16 hours. The AF precipitated as a red, gel-like precipitate which we collected by centrifugation at 10,000 x g. We washed this pellet 3 times in Tris-buffered MBL-seawater (422 mM NaCl, 9.4 mM KCl, 9 mM CaCl_2_, 49.4 mM MgCl_2_, 28 mM MgSO_4_ • 7H_2_0, 0.85 mM NaHCO_3_, 50 mM Tris, pH 7.2) for proteomic analysis as the “Crude” AF sample.

Following Varner and colleagues (12) we further purified the crude AF fraction by resuspending the pellet in CMFSW, passing it through a 0.22 µm filter, centrifuging at 20,000 x g to remove red contaminating pigments and membranes, then precipitating it in 20 mM CaCl_2_ overnight at 4 ºC. The pellet that formed was white and fluffy in appearance. We again washed this pellet in Tris-buffered MBL-seawater prior to proteomic analysis. This constituted the “Filtered” sample.

As the final purification step, we re-solubilized the Filtered AF fraction in CMFSW + 1mM EDTA (it remained insoluble in CMFSW alone). We then concentrated the sample as much as possible without causing the AF to precipitate in an Amicon-Ultra 10 column (EMD Millipore), and loaded it onto a Sephacryl S-500 HR column (Cytiva). The AF eluted as a single broad peak which we concentrated in an Amicon-Ultra 10 column and analyzed by proteomics as the “SEC” sample.

### Atomic Force Microscopy Imaging and Instrumental procedure

Atomic Force Microscopy (AFM) imaging was acquired under ambient conditions in the tapping mode of operation using standard monolithic Si cantilevers Tap300AI (NanoAndMore) on a commercial instrument Multimode Nanoscope IIIa (Veeco). The aggregation factor complexes were immobilized via physisorption from solution on muscovite mica surfaces (Plano), which were previously gas phase-silanized with aminopropyltriethoxysilane (Sigma) in a desiccator. The immobilization of the AFs from solution (50 *µ*L of typically 0.5 mg/mL) for 15 min at room temperature was subsequently followed by a washing step with Milli-Q water as well as dried under N_2_ flow.

### Preparation of Samples for Proteomic Analysis

We lyophilized the samples and treated the pellets with freshly prepared hydroxylamine buffer (1 M NH_2_OH-HCl, 4.5 M guanidine-HCl, 0.2 M K_2_CO_3_, pH adjusted to 9.0 with NaOH). We briefly vortexed the samples and then incubated them at 45 ºC for 6 hours. Due to pressure build-up during incubation, we fastened the tubes shut during incubation. After incubation, we spun the samples for 15 min at 18,000 x g, removed the supernatant, and stored it at -20 ºC until further proteolytic digestion with trypsin.

We digested the samples following the filter-aided sample preparation (FASP) protocol, using a 10 kDa molecular weight cutoff filter. Briefly, we mixed 50 µL of samples in the filter unit with 8 M urea and 100 mM ammonium bicarbonate (AB), pH 8.0, and then centrifuged the mixture at 14,000 x g for 15 minutes. We reduced the proteins with 10 mM DTT for 30 min at room temperature, centrifuged them, and alkylated them with 55 mM iodoacetamide for 30 min at room temperature in the dark. After centrifugation, we washed the samples three times with urea solution and three times with 50 mM AB, pH 8.0. We carried out protein digestion using sequencing grade modified Trypsin (Promega) at a 1/50 protease/protein (wt/wt) ratio, incubating at 37 °C overnight. Finally, we recovered the peptides from the filter using 50 mM AB.

### Proteomics LC-MS/MS measurements

We loaded 20 µL of each sample onto individual Evotips (Evosep, Odense Denmark) for desalting, washed them with 20 µL of 0.1% FA, and then added 100 µL of storage solvent (0.1% FA) to keep the Evotips wet until analysis. The Evosep One system (Evosep) was used to separate peptides on a Pepsep column (150 um inner diameter, 15 cm) packed with ReproSil C18 1.9 um, 120A resin using pre-set 15 samples per day gradient. We coupled the system to the timsTOF Pro mass spectrometer (Bruker Daltonics in Bremen, Germany) via its nano-electrospray ion source, Captive Spray.

We operated the mass spectrometer in PASEF mode, setting the ramp time to 100 ms and acquiring 10 PASEF MS/MS scans per topN acquisition cycle. We recorded MS and MS/MS spectra from m/z 100 to 1700 and scanned the ion mobility from 0.7 to 1.50 Vs/cm^2^. We isolated precursors for data-dependent acquisition within *±* 1 Th and fragmented them with an ion mobility-dependent collision energy, linearly increasing it from 20 to 59 eV in positive mode. We repeatedly scheduled low-abundance precursor ions that had an intensity above a threshold of 500 counts but below a target value of 20,000 counts, and dynamically excluded them for 0.4 minutes.

### Proteomics Database Searching and Protein Identification

Mass spectrometry raw timsTOF ddaPASEF .d files were processed using the default LFQ-MBR workflow within Fragpipe v21.1 (79). Files were converted to mzBIN format and then searched using MSFragger v4.0 (80) against the predicted *C. prolifera* proteome including known contaminants and the reversed protein sequences. The default search parameters were as followed: strict tryptic digestion with a maximum of 2 missed cleavages, peptide length = 7-50, peptide tolerance = 20 ppm; MS/MS tolerance = 10 ppm; topN peaks = 150; fixed modifications = carbamidomethyl on cysteine; variable modifications = acetylation on protein N termini and oxidation of methionine. Peptide Spectrum Matches (PSMs) validation was performed by philosopher version 5.1.0 and the false discovery rate was fixed at 1% at the PSMs, peptides and proteins level. Label free quantification on the MS1 level using unique as well as razor peptides was conducted by Ionquant (81) using default settings.

The raw output files of FragPipe (protein.tsv files) were processed using the R and python programming languages. Contaminants and reverse proteins were filtered out and only proteins that were quantified with at least 2 razor peptides (Razor.Peptides >= 2) were considered for the analysis. In order to compare relative protein proportions within a sample (crude, filtered or SEC), individual protein intensities (based on razor peptides) were divided by the sum of all protein intensities.

### Annotation and Analysis of Top Proteomic Hits

In addition to the annotation tools built-in to the TransPi pipeline, we annotated the *C. prolifera* proteome using EggNOG Mapper v2 (84). We then manually examined the most abundant hits in our proteomic analysis by BLAST search against the NCBI nr and Eukprot3 databases, and by using Interpro to predict the presence of signal peptides, conserved domains, and disordered regions. settings.

We used the published Wreath domain HMM (33) to search for Wreath domain-containing proteins in *C. prolifera* using HMMer (84, 90), and used Alphafold2 (91) as implemented by Colabfold (38) to predict the tertiary structure of Wreath domain-containing proteins represented in our proteomics datasets. The MAFp3 Wreath domain structural model was then used as an input into default Foldseek (41) (v8-ef4e960) to search for related structures in other species. settings. We identified the candidate AF-interacting region by BLAST search of *C. prolifera* MAFAP1 against the *C. prolifera* predicted proteome. We then used an alignment of the conserved C-terminal region of top BLAST hits as an input into HMMer to create an HMM for the AF-interacting region. We used the AF-interacting region HMM to search additional sponge genomes and transcriptomes for homologous proteins. We used ESMFold (82) as implemented in the Foldseek web server to predict the tertiary structure of the MAFAP1 AF-interacting region, and Foldseek to search for structurally similar proteins in other species. settings.

Visualization and superposition of *C. prolifera* wreath domain beta-sandwich and H. sapiens Mucin-5AC VWFD1 domain (UniprotID: P98088, aa 79-249) was done in Pymol (v2.3.5) using the “super” command. Structural superposition of ESMFold prediction of region 1 and 2 of MAFAP1 C-terminal domain and NCAM1a Ig-like- and Fn3 domain (UniprotID: P16170) was done using the “cealign” command.

Transmembrane region prediction was performed using the DeepTMHMM web server (v.1.0.24) (49). GPI-anchor prediction was performed using the NetGPI 1.1 web server (50).

### Analysis of MAFAP1-like proteins in related sponges

We next examined the prevalence of AF-associated proteins using published sequence data from additional sponge species. First we searched published datasets for *Ephydatia muelleri* (92), *Cladhoriza sp*. (54), and *Phorbas areolatus* (57). We then downloaded raw transcriptome reads from NCBI (SRA) for available poecilosclerid sponges: *Asbestopluma hypogea* (ER216190), *Crella elegans* (SRR648671), *Isodictya* (SRR6202908-12), *Kirkpatrickia variolosa* (SRR1916957), *Latrunculia apicalis* (SRR1915755), *Mycale grandis* (SRR3334580, SRR3339390, SRR3339394), *Mycale phyllophila* (SRR1711043, SRR2394941, SRR2402290), and *Tedania anhelans* (SRR3708911). Raw Illumina reads were error-corrected using Rcorrector (93) using a k-mer length of 31. The corrected reads were then quality-trimmed using Trimmomatic via Trinity v.2.4.0 with the default settings (86). The trimmed reads were combined within species and assembled with Trinity v.2.4.0 using the default settings. Raw 454 pyrosequencing reads were assembled using the default settings in MIRA v.4.0 (86, 87). We then translated the assemblies to peptide predictions using TransDecoder (https://github.com/TransDecoder/TransDecoder). The peptide predictions were then clustered to 99% similarity using CD-HIT (94) to reduce redundancy from splice isoforms and in-paralogues. To reduce potential protist and prokaryote contaminants, we filtered the peptide assemblies using Alien Indexing (https://github.com/josephryan/alien_index) using the provided metazoan and non-metazoan representative datasets for the BLAST searches, with peptide models from the *Tethya wilhelma* genome (52) supplementing the metazoan dataset. We then searched the decontaminated and translated poecilosclerid assemblies, as well as genome models from *T. wilhelma* and *Amphimedon queenslandica*, for MAFP1-related proteins using the *C. prolifera* candidates as queries in BLAST searches (e-values 1e-50 and 1e-150). We also searched the additional sponges using a Hidden Markov Model of the C-termini with hmmsearch in HMMER3 (v3.1b2) (89).

## Supplementary Files and Figures

**Supplementary File 1**: *C. prolifera* wreath domain containing proteins.

**Supplementary File 2**: *C. prolifera* MAFp3 wreath domain Colabfold structural prediction.

**Supplementary File 3**: *C. prolifera* MAFAP1 C-terminus Colabfold structural prediction.

**Supplementary File 4**: *C. prolifera* MAFAP1 C-terminus HMM profile.

**Supplementary File 5**: *Tedania anhelans* homologs of AF-interacting proteins.

**Supplement Figure 1.**
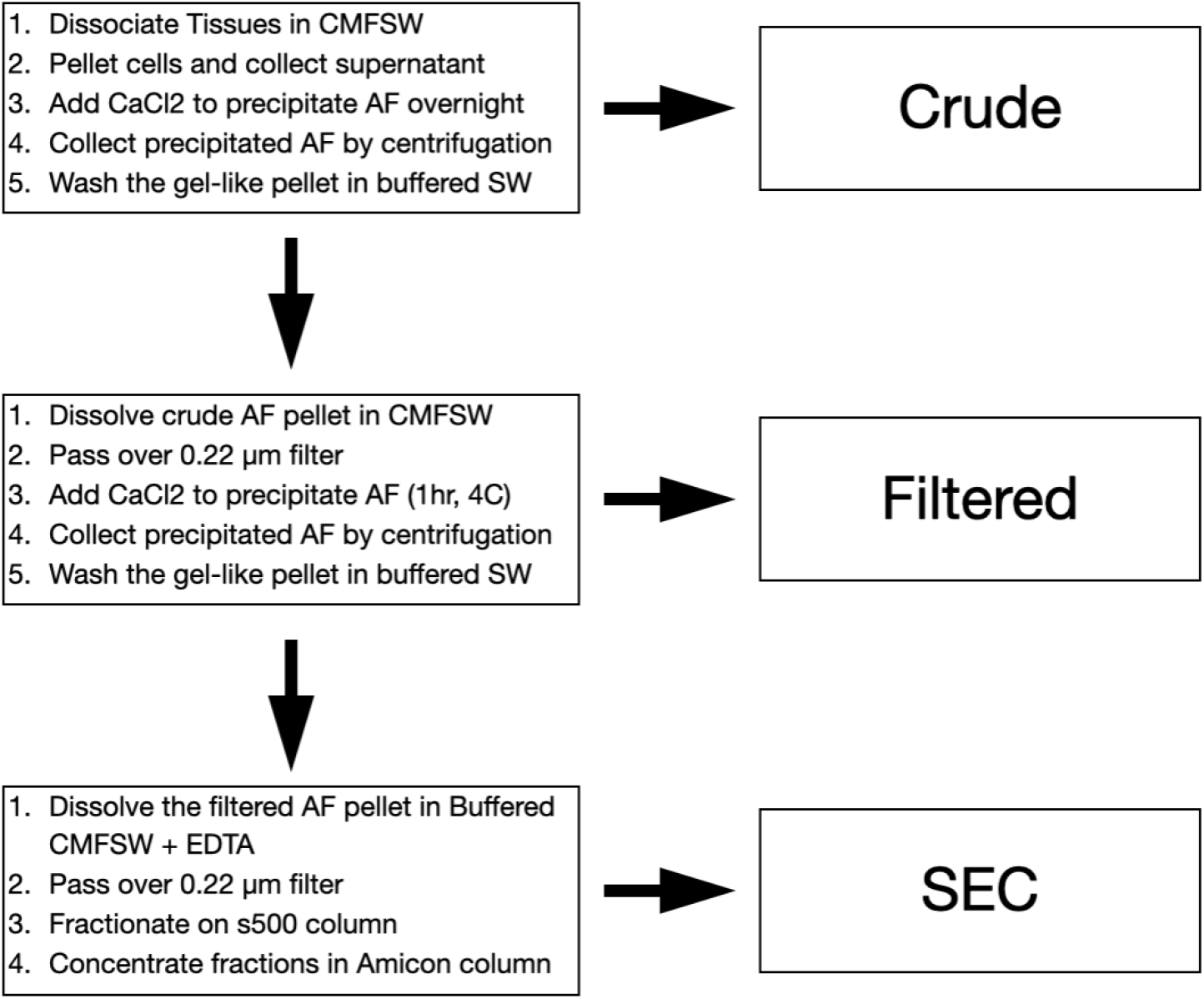
Sample preparation for proteomics analysis. Workflow used to prepare Crude, Filtered, and SEC aggregation factor samples for proteomics analyses.

**Supplement Figure 2.**
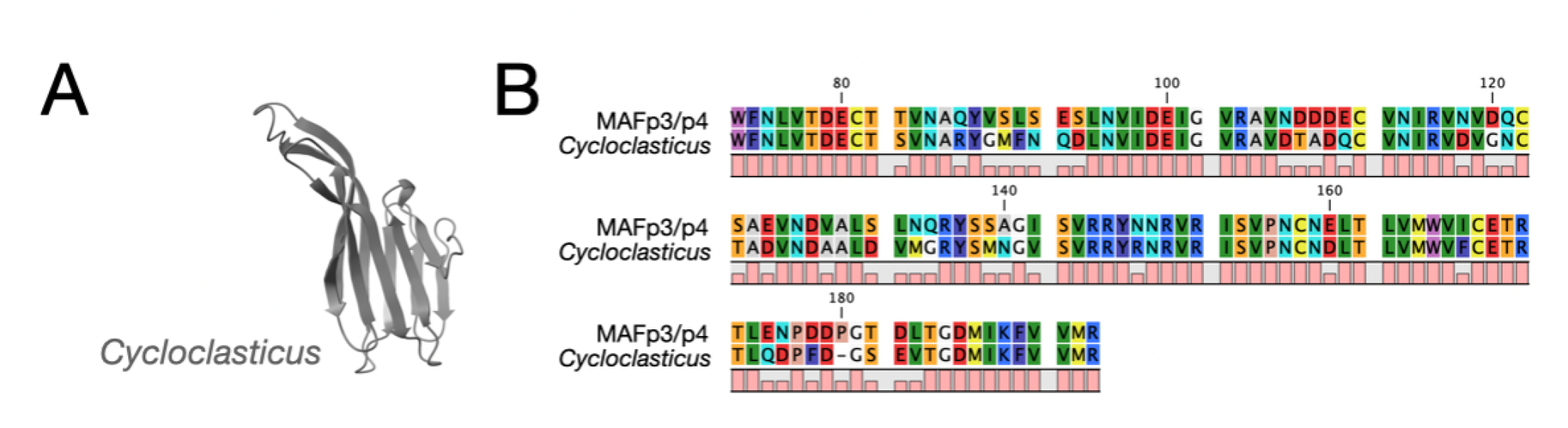
Comparative analysis of Wreath domain in other species. **(A)** Foldseek search using the *C. prolifera* MAFp3 wreath domain as a query revealed structural similarity to a protein domain predicted from the bacterium *Cycloclasticus sp*. (A0A1X1QQN9), which was isolated as a symbiont of a poecilosclerid sponge. **(B)** Alignment to the *C. prolifera* MAFp3 wreath domain reveals extensive sequence conservation.

**Supplement Figure 3.**
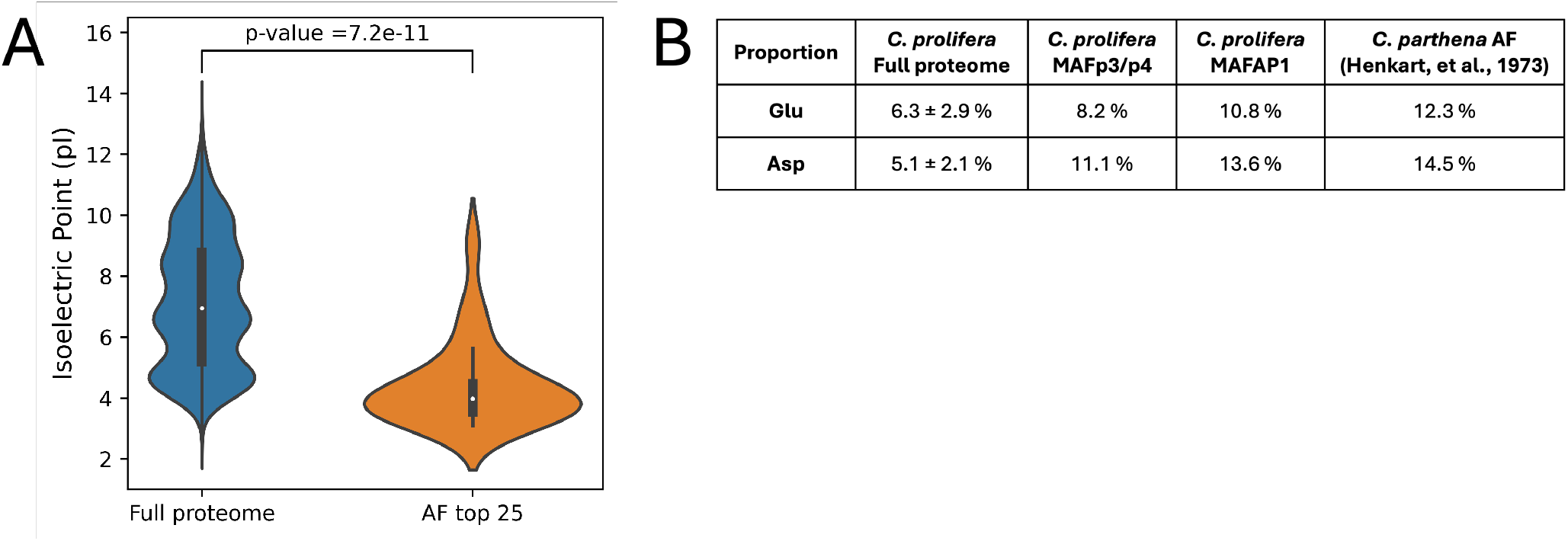
AF-associated proteins are predicted to be negatively charged at neutral pH. **(A)** Distribution of isoelectric points from *C. prolifera* full proteome vs. top 25 most abundant proteins in the AF sample. P-value was calculated using the Wilcoxon rank sum test. **(B)** Proportion of Glu and Asp residues across different samples.

