## Supplemental File 5 for "Proteomic analysis of the Aggregation Factor from the sponge *Clathria (Microciona) prolifera* suggests an ancient protein domain toolkit for allorecognition in animals"

**Frizzled Related proteins with a conserved AF-interacting C-terminus**

>Tedania_anhelans_SRR3708911__TRINITY_DN13195_c2_g1_TRINITY_DN13195_c2_g1_i8_g_89973_m_89973
MIGHLRIISAIKGWLGIILFISPLSLQQGVRIVSPNEPTYHCEELTVITWCSYIYPNASFPNYRGHEDQFSANRELLNFTPLVQSVCSNAIVHFLCSIYAPFCQPDLPNHRIRPCRELCDYVRSTCEEDLQNFNLEWPPHLDCDNFMPNSATNLDFCPENLHLLRIPAQIPTFTSPSATSPATEEGGASTTTTQVQPEGTTTEQSTTLTQTTTSDNTTTTTQDQPEGTTTQESVTVTTIQPPTTRPMPSLYFELNGQTYPSGSKISITDIGVANPNNLLDTASALVCVTSEVNTQCCRRRDGGNVGEWFLPDGSIVPRLRNSPNGDFTRSGFRHHVRLNRRNDAMAPLGTYTCLVPDERDGSVLHTATLELEVPGILPREVLCFCSVCVWRAPEGSLDRHRGYDIRFSGELVIGKGNDEFFHVVAGDERITALGLQEGVTVEVRVRSEGETVNPWSGPVSLCSNE*

>Tedania_anhelans_SRR3708911__TRINITY_DN13195_c2_g1_TRINITY_DN13195_c2_g1_i5_g_89969_m_89969
MIGHLRIISAIKGWLGIILFISPLSLQQGVRIVSPNEPTYHCEELTVITWCSYIYPNASFPNYRGHEDQFSANRELLNFTPLVQSVCSNAIVHFLCSIYAPFCQPDLPNHRIRPCRELCDYVRSTCEEDLQNFNLEWPPHLDCDNFMPNSATNLDFCPENLHLLRIPAQIPTFTSPSATSPATEEGGASTTTTQVQPEGTTTEQSTTLTQTTTSDNTTTTTQDQPEGTTTQESVTVTTIQPPTTRPMPSLYFELNGQTYPSGSKISITDIGVANPNNLLDAASALACVTSEVNTQCCRRRDGGNVGEWFFPDGSIIPRLRNSPNRDFTRSGFMHQVRLNRKNDAMAPLGTYTCRVPDAENSAVVHTASITLVAPGILPREVLCFCSVCVWRAPEGSLDQHRGYDIRFAGELVISKENDEFFHVVAGDERIAALEPQEEATVEVRVSSEGEAANPWSDPVSLCSNNNLRM*

Frizzled domain
AF-interacting region

**Calx-beta-containing proteins with a conserved AF-interacting C-terminus**

>Tedania_anhelans_SRR3708911__TRINITY_DN13195_c2_g1_TRINITY_DN13195_c2_g1_i11_g_89979_m_89979
TVTESAGFQEVCVQVFNPPSNEESVFDIFLVHKTTTGSAAEFLDFSPVFDLQLFPPSLPEEGVTRRGCFNVFIADDELCENTESFTISLELDTFTSQPGVRVDPSITEIFIVDDDVVIGFTDGPYAASENDGFAAISFGVIDGTLQSALVELSFTGGTALIGEDFILDAESTYTLTANNPTVQVQVPLIDDAVFEIAENLTAHLSLTDEARPCVTISPDSAEITIEDNDILTFGFTSDVYEFGEDSGANLIFVSLLSGNPGEFSVVLTAATDRNSTNATATVGLDYNEVNATQLEFSSSLSNTASFNVTIICDFLTEAREFFEVKITNILVRKSLGPQLYRRIVKSKWGRRILFVVKTAQIIIRNINKVATSTEGMLTTQPGVIGTTIQPPTSSDNTTTTQDQPEGTTTQESVTVTTIQPPTTTSDNTTATSQNQPEETTTQESVTVTIQPPTTRPMPSLYFELNGQTYPSGSKILITDIRVANPNNLLDAASALVCVTSEVNTQCCRRSDGGNVGEWFFPNGSIVLRSRNSPNRDFTRSGFRHQVRLNRKNDAMAPLGTYTCLVPDESDSAVLHTASVTLELKVPGILPHEVLCFCSVCVWRAPEGSLDRHRGYDIRFSGELVIGKENDEFFHVVAGDERIAALGPQEGVTVEVRVRSEGETVNPWSDPVSLCSNE*

Calx-beta domain
AF-interacting region
